# Markov models of the apo-MDM2 lid region reveal diffuse yet two-state binding dynamics and receptor poses for computational docking

**DOI:** 10.1101/053603

**Authors:** Sudipto Mukherjee, George A. Pantelopulos, Vincent A. Voelz

## Abstract

MDM2 is a negative regulator of p53 activity and an important target for cancer therapeutics. The N-terminal lid region of MDM2 modulates interactions with p53 via competition for its binding cleft, exchanging slowly between docked and undocked conformations in the absence of p53. To better understand these dynamics, we constructed Markov State Models (MSMs) from large collections of unbiased simulation trajectories of *apo*-MDM2, and find strong evidence for diffuse, yet two-state folding and binding of the N-terminal region to the p53 receptor site. The MSM also identifies *holo*-like receptor conformations highly suitable for computational docking, despite initiating trajectories from closed-cleft receptor structures unsuitable for docking. Fixed-anchor docking studies using a test set of high-affinity small molecules and peptides show simulated receptor ensembles achieve docking successes comparable to cross-docking studies using crystal structures of receptors bound by alternative ligands. For p53, the best-scoring receptor structures have the N-terminal region lid region bound in a helical conformation mimicking the bound structure of p53, suggesting lid region association induces receptor conformations suitable for binding. These results suggest that MD+MSM approaches can sample binding-competent receptor conformations suitable for computational peptidomimetic design, and that inclusion of disordered regions may be essential to capturing the correct receptor dynamics.

## Introduction

Under normal cellular conditions, the tumor suppressor protein p53 is kept at a low basal level in part due to downregulation by MDM2 (mouse double minute 2 homolog), an E3 ubiquitin ligase that recruits p53 for degradation via direct interaction with the p53 transactivation domain (TAD). Since many tumor cells still retain wild-type p53, a promising ave-nue of cancer treatment is to restore p53 activity by blocking the MDM2-p53 interaction with high-affinity MDM2-binding ligands.

A high-resolution x-ray crystal structure of p53 TAD bound to MDM2 in a helical conformation has been available for some time, and has spurred widespread effort towards developing inhibitors that potently disrupt p53-MDM2 binding.^1^ In addition to small molecules.^2,3^ peptidomimetics have been designed to mimic the p53 helix, such as stapled peptides,^4^ beta-peptides,^5^spiroligomers,^6^ high-affinity D-peptides,^7^^−^^9^ arylamides, terphenyls, hydrogen-bond surrogates^10^ and oligooxopiperazines,^11^many of which were developed as a result of-or in concert with-computational modeling and design.^3,11−^^16^

Aside from its therapeutic interest, the p53-MDM2 interaction has served as a valuable model system for understanding protein-protein interactions, especially for intrinsically disordered proteins such as the p53 TAD that fold upon binding.^17^ Underscoring the importance of this work is recent evidence that residual helicity in the p53 TAD directly alters cell signaling in vivo.^18^ Similarly, consideration of intrinsic disorder is important to understanding MDM2, as it contains an unstructured N-terminal lid region (residues 1-25) which competes with p53 for the binding cleft. In the absence of p53, quantitative NMR spectroscopy has shown transient structuring and binding of the lid region to the p53 cleft on slow (>10 ms) exchange timescales, consistent with the structuring of a short, well-ordered helix in residues 19-24 (Figure 1).^19^ Recent NMR and X-ray co-crystal structures have revealed that small-molecule inhibitors can induce structuring of the lid region through specific favorable interactions,^20^ suggesting that computational prediction lid region structure and dynamics could be very useful for computational design.

**Figure 1.**
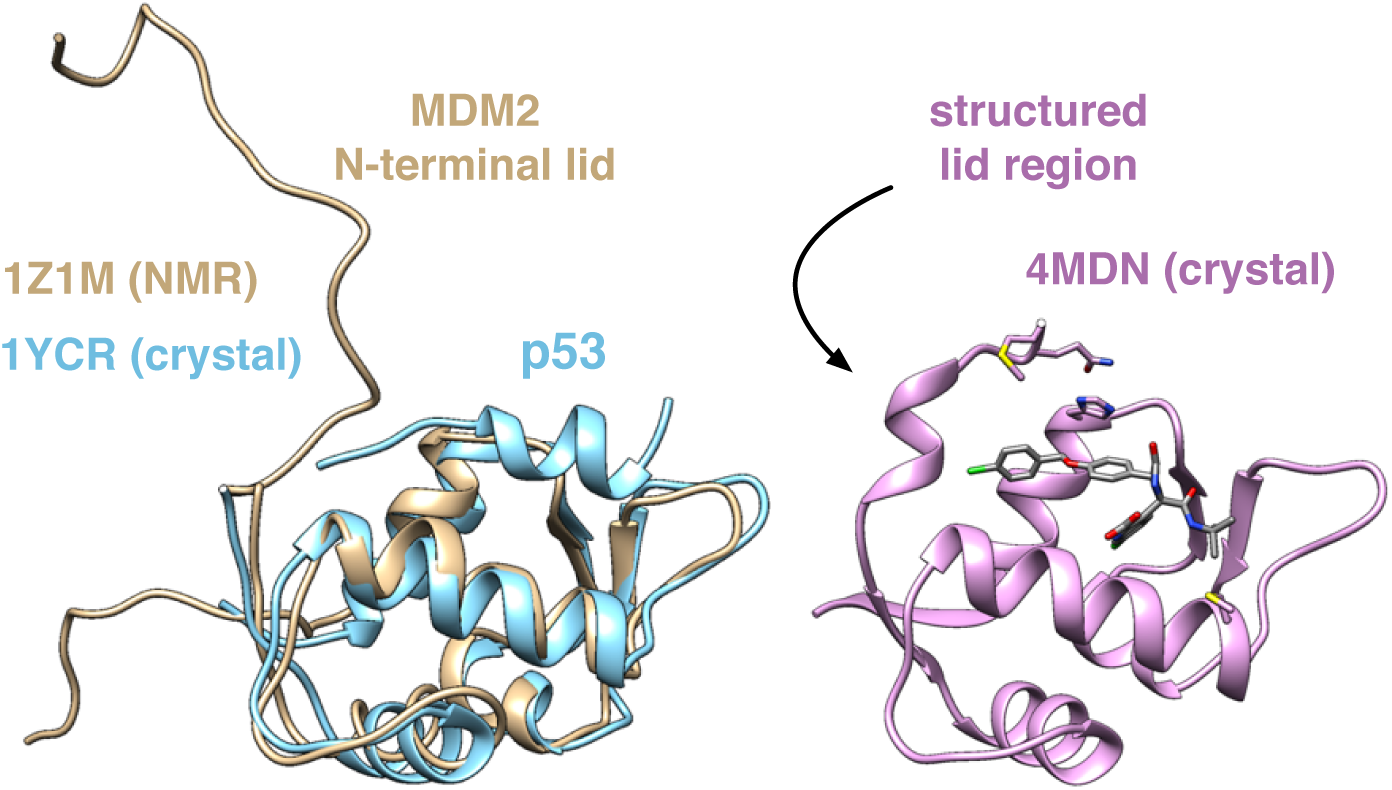
*Apo* and *holo* structures of MDM2. (left) The *apo* form of MDM2 (tan) has an unstructured N-terminal lid (residues 1-25) that associates with the cleft. The binding of p53 displaces the lid region from the binding cleft. Quantitative NMR of *apo*-MDM2^19^ has shown that a portion of the lid region (residues 19-24) slowly converts between an unstructured and a structured state. A recent co-crystal structure^20^ with a small-molecule inhibitor (pink) shows a structured form of the lid region.

Here, to better understand the structure and dynamics of the N-terminal lid region of *apo*-MDM2, we perform extensive simulation studies to characterize the mechanism of association with the p53 binding cleft, and explore the possible role of such computational studies in drug discovery. From many independent trajectories of MDM2 starting from the *apo* state obtained by parallel distributed simulation, we construct a Markov State Model (MSM) of N-terminus dynamics that predicts two-state binding to the p53 cleft, in agreement with experimental findings. We then explore the utility of the MSM for *in silico* drug discovery by performing computational docking studies to kinetic metastable states of the MDM2 receptor. Remarkably, our findings suggest that the ensemble of metastable receptor conformations identified in the MSM can be used to achieve docking results similar to or better than cross-docking studies of crystal structures, and moreover, that inclusion of the N-terminus is essential in selecting open-cleft receptor conformations suitable for docking.

## Results

### Markov State Model (MSM) analysis of simulated *apo*-MDM2 dynamics predicts two-state binding of the lid region to the p53 cleft

MSMs describe conformational dynamics as a network of transitions between kinetically metastable states.^21^ To construct an MSM of N-terminal dynamics from simulation data, trajectory snapshots are first assigned to metastable conformational states. To identify these metastable states, we first used tlCA^22,23^ to find a low-dimensional subspace reflecting the slowest conformational motions of the N-terminal region (residues 1-25) and residues in the binding site. Projections to the two largest components (tIC1 and tIC2) were subsequently used for conformational clustering into 2000 metastable microstates, and for visualizing the folding/binding landscape.

Next, observed transitions between states are used to infer a transition matrix **T**^(*τ*)^, whose elements *T*_*ji*_, contain the probability of transitioning from state *i*to state *j*within time *τ*. The right (*ϕ*_*n*_) and left (*ψ*_*n*_) eigenvectors of the transition matrix yield a complete description of state population dynamics, via the chemical master equation, *d***p**/*dt* = **Kp**, where **T** = exp(*τ***K**), whose solution is

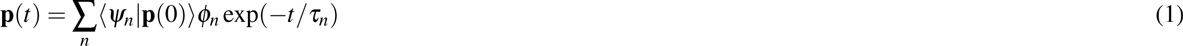

Here, p(0) is a vector of initial state populations at time *t* = 0, and the implied timescales *τ*_*n*_ = —*τ*/1nµ_*n*_ associated with each eigenmode n are related to the eigenvalues *µ*_*n*_ of T. We define the sign structure of each eigenvector such that (*ψ*_*n*_| l) is positive, so that dynamics (starting from a hypothetical uniform distribution) can be described as a superposition of positive-amplitude eigenmodes *ϕ*_*n*_ each decaying at time scale *τ*_*n*_. The stationary eigenvector (i.e. the equilibrium state populations) is *ϕ*_0_, for which *τ*_*n*_ = ∞. The resulting MSM clearly shows a two-state mechanism for lid region binding to the cleft. The sign structure of the slowest relaxation eigenmode *ϕ*_1_ shows population flux from unbound to bound states of the lid regions, indicated by two diffuse basins aligned with tIC_1_, the degree of freedom representing the slowest conformational motions (Figure 2a). Interestingly, compared to the tICA landscapes reported for many protein folding systems,^24^^−^^26^ the lid landscape is remarkably diffuse, even in the secondary eigenmodes (Figure S4), reflecting the lack of residual structure. Similar landscapes have been found in other MSM studies of disordered proteins.^27,28^ Implied timescales computed at lag times ranging from 100 ps to 100 ns show a clear gap between the slowest and next-slowest implied timescale, indicative of apparent two-state dynamics (Figure 2b). The slowest implied timescale is close to 1 *µ*s, comparable to the molecular on-rate of a peptide at high effective concentration. This timescale is over four orders of magnitude faster than the slow (> 10 ms) conformational exchange of residues 19-24 reported by Showalter et al., which suggests that our simulation trajectories, each shorter than 1 *μ*s, do not capture rare unstructuring events expected this region. Nevertheless, the simulations show good agreement with experimentally measured chemical shifts in this regions for the apo state, which is estimated to have ~90% of the lid population in an associated state.

**Figure 2.**
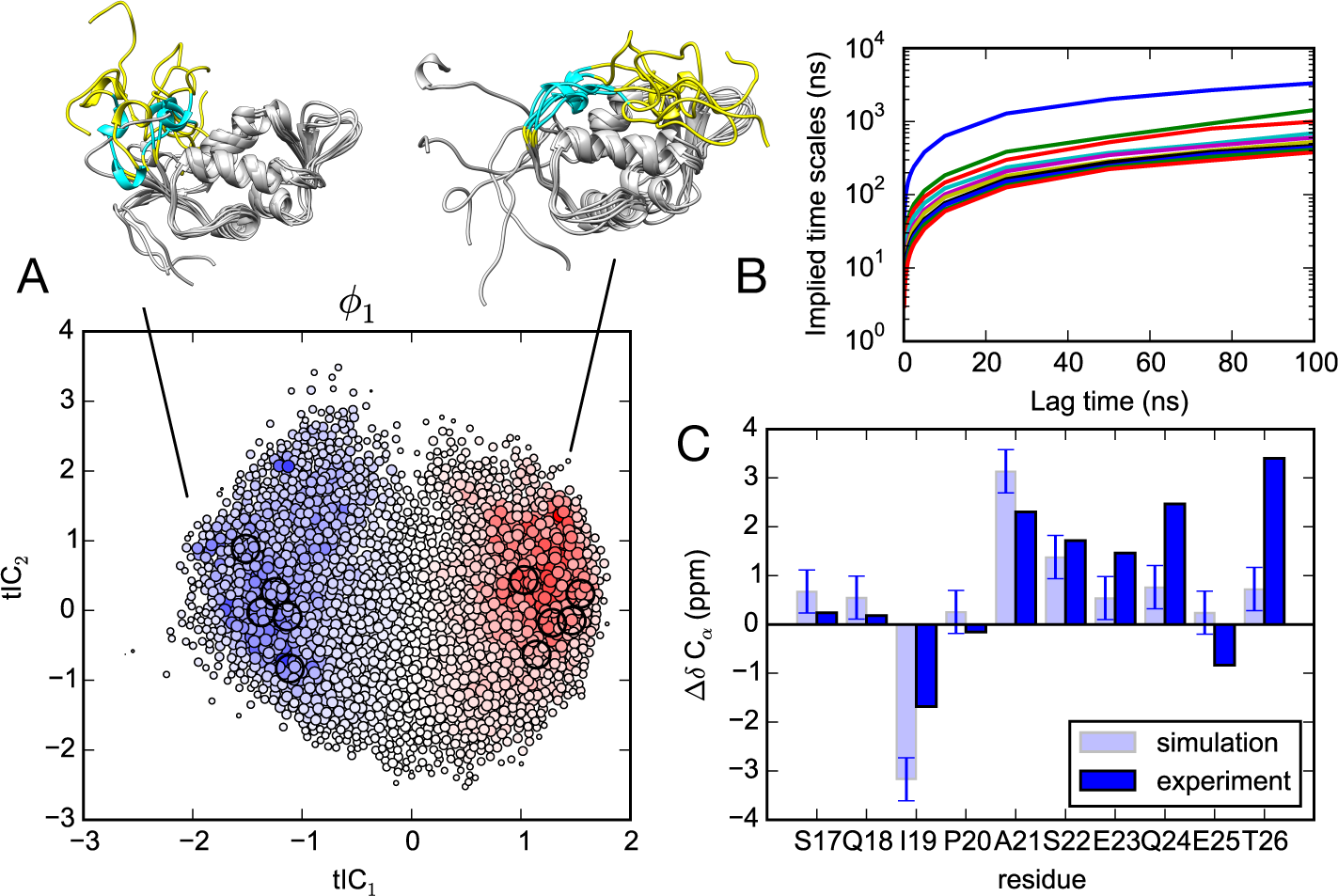
(A) Projection of the 2000 MSM microstates (filled circles) to tIC_1_ and tIC_2_ coordinates. The size of each circle is proportional to the equilibrium population, and is colored according to the slowest relaxation eigenmode, *ϕ*_1_. Population flux along this mode is from blue to red, representing a transition from unbound to bound states of the lid region, which we visualize using five representative structures from each basin. (B) Implied timescales versus MSM lag time show a clear gap indicating apparent two-state kinetics. (C) Simulation predictions of C_*α*_ chemical shift deviations from random-coil for the lid region (residues 17-26, cyan ribbon in panel A) calculated by SHIFTX2^29^ agree with experimental values^19^.

The next-slowest eigenmode relaxation, *ϕ*2, reflects conformational dynamics of the lid region along the tIC_2_ component, and it is similarly diffuse (Figure S4). To gain structural insight into these motions, we performed secondary structure analysis and Bayes Factor analysis^25^ of interresidue contacts formed along different quadrants of the tICA projection (SI Text, Figures S4 and S5). While the slowest relaxation (along tIC1) corresponds to disassociation of the N-terminus from the C-terminus, structuring of the lid region into a helix, and association with the binding cleft helix α2, the next-slowest relaxation (along tIC_2_) largely reflects an increase in average self-association of the lid region, with an increase in sheet structure.

### Computational docking of known MDM2 ligands to simulated receptor ensembles achieves success comparable to crystal structure cross-docking

Virtual screening studies rely heavily on the availability of high-resolution crystal structures. Since the MDM2 trajectories were initiated from an *apo*NMR structure (PDB: 1Z1M) with a closed binding cleft unsuitable for computational docking, our work presents an excellent opportunity to test how successfully an MD+MSM approach can be used as a refinement procedure to achieve high-quality receptor structures for docking.

To evaluate the quality of simulated receptor structures, we used the DOCK6 algorithm to perform computational docking of a test set of 10 ligands to the 2000 MSM microstate structures (with the lid region removed). Our test set consisted of eight small-molecule ligands and two peptide ligands, all with high-resolution crystal structures (Table 1). The small-molecule ligands include, among others, the best-in-class inhibitor nutlin, and similar compounds. The peptide ligands include the native p53 fragment,^30^ and a high-affinity designed inhibitor sequence, PMI N8A.^31^ Several modifications were made to standard docking procedures to facilitate the efficient docking of peptide sequences, most notably: fixing backbone atoms in their helical conformation via an artificial cyclization bond between terminal alpha-carbons, while retaining side chain rotamer search (see Methods).

**Table 1.**
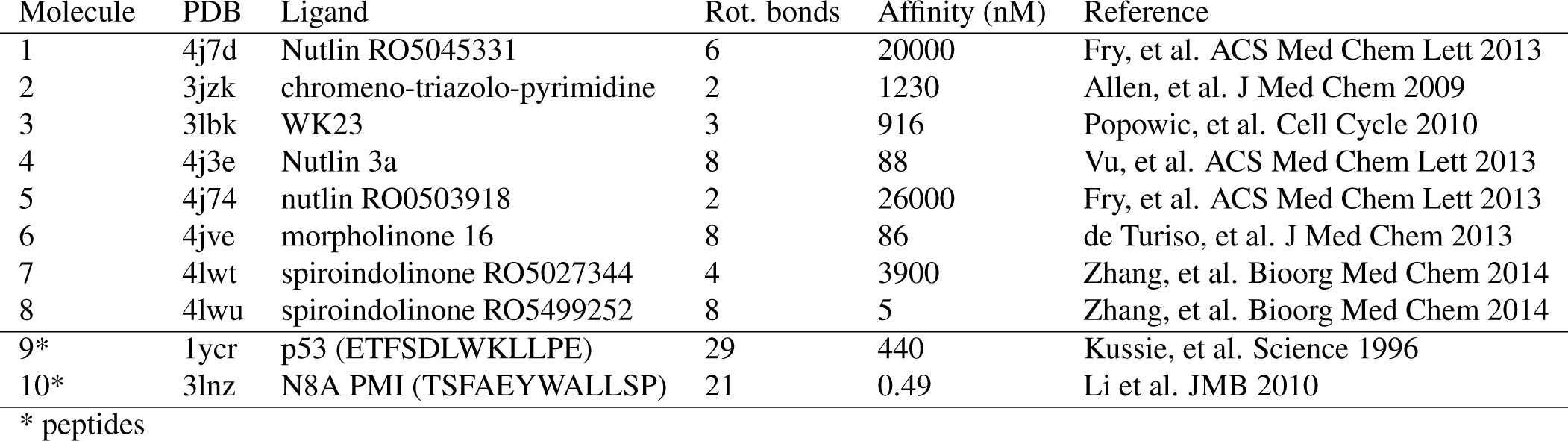
Test set of small-molecule and peptide ligands of MDM2 used for computational docking studies.

| Molecule | PDB | Ligand | Rot. bonds | Affinity (nM) | Reference |
| --- | --- | --- | --- | --- | --- |
| 1 | 4j7d | Nutlin RO5045331 | 6 | 20000 | Fry, et al. ACS Med Chem Lett 2013 |
| 2 | 3jzk | chromeno-triazolo-pyrimidine | 2 | 1230 | Allen, et al. J Med Chem 2009 |
| 3 | 3lbk | WK23 | 3 | 916 | Popowic, et al. Cell Cycle 2010 |
| 4 | 4j3e | Nutlin 3a | 8 | 88 | Vu, et al. ACS Med Chem Lett 2013 |
| 5 | 4j74 | nutlin RO0503918 | 2 | 26000 | Fry, et al. ACS Med Chem Lett 2013 |
| 6 | 4jve | morpholinone 16 | 8 | 86 | de Turiso, et al. J Med Chem 2013 |
| 7 | 4lwt | spiroindolinone RO5027344 | 4 | 3900 | Zhang, et al. Bioorg Med Chem 2014 |
| 8 | 4lwu | spiroindolinone RO5499252 | 8 | 5 | Zhang, et al. Bioorg Med Chem 2014 |
| 9* | 1ycr | p53 (ETFSDLWKLLPE) | 29 | 440 | Kussie, et al. Science 1996 |
| 10* | 3lnz | N8A PMI (TSFAEYWALLSP) | 21 | 0.49 | Li et al. JMB 2010 |

\* peptides

To establish the baseline accuracy of the DOCK algorithm for this system, ligands were re-docked to their own co-crystal structures, and cross-docked to all other receptor structures in the test set (Figure 3).

**Figure 3.**
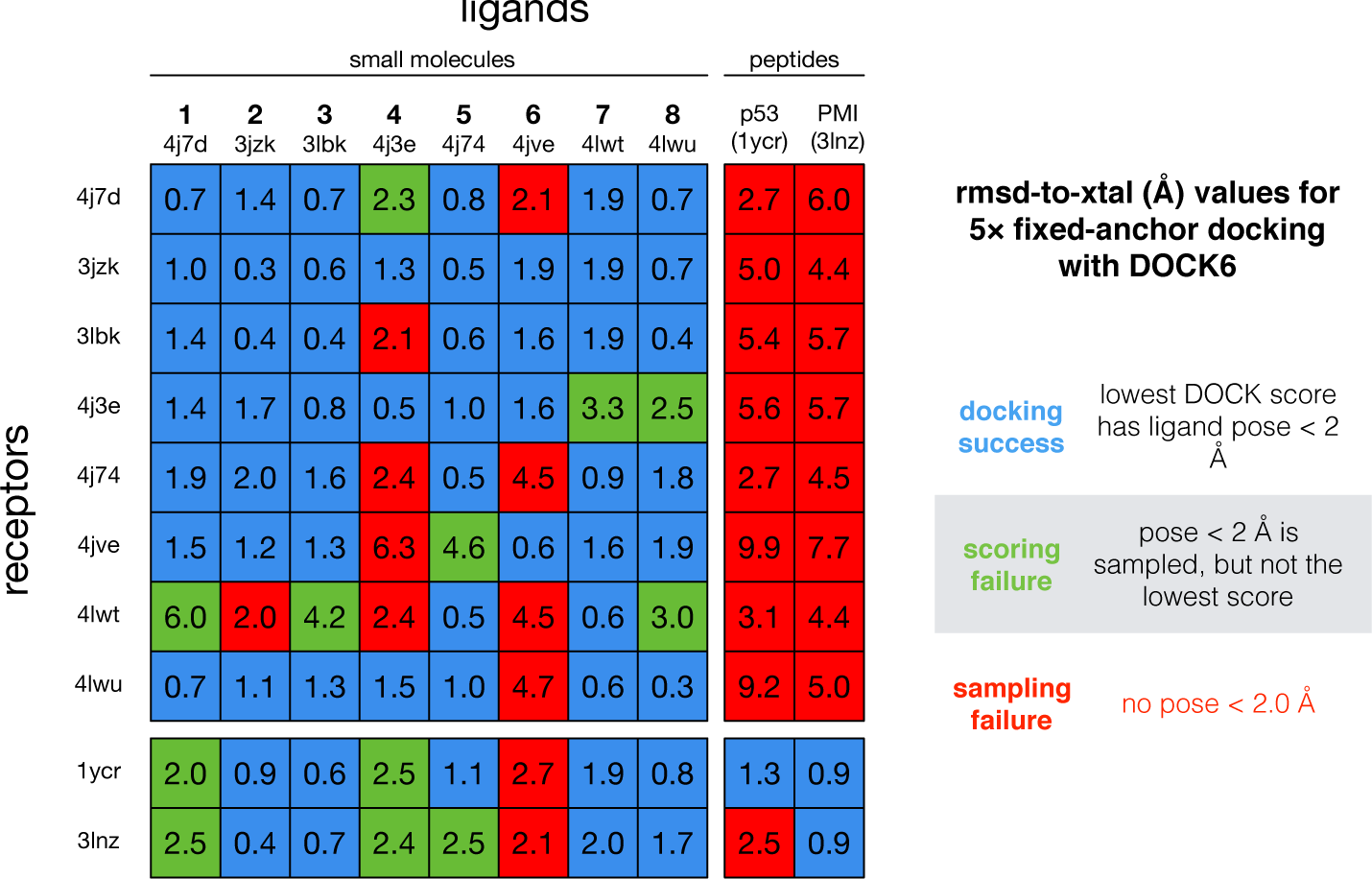
Self-docking and cross-docking results for a test set of 10 MDM2 ligands with available co-crystal structures (listed by PDB ID). Values shown are the rmsd (in Å) of the best-scoring docked ligand pose to the crystal ligand pose. Docking successes are shown in blue, scoring failures are shown in green, and sampling failures are shown in red.

In all cases, the best re-docking scores corresponded to a correctly docked pose, which we define as having an rmsd of 2.0 Å or less to the crystal pose, thus validating the accuracy of DOCK. Cross-docking results show the inherent variability of docking to a target receptor structure, and show that some MDM2 receptor crystal structures are more likely to produce false positives or outright failures when non-native ligands are docked. Cross-docking is the least successful for small-molecule docking to peptide-bound receptor structures, and vice versa. We also cross-docked all the ligands in the test set to the *apo*-MDM2 receptor structure (PDB:1Z1M, with the lid region removed), which confirmed its unsuitability for docking; best-scoring poses for all ligands showed rmsd values > 5.4 Å.

By comparison, computational docking to the ensemble of 2000 MSM microstates is much more successful. Plots of the DOCK score versus ligand pose rmsd show a funnel-like correlation, indicating that low scores indeed predict good ligand poses (Figure 4). Because of this, a significant enrichment in correct docking predic-tions is achieved. If only the five best-scoring receptor poses were considered (the top 0.25%), half of the ligands would be correctly docked; 80% are correctly docked if only the top 20 (1%) receptor poses are considered.

**Figure 4.**
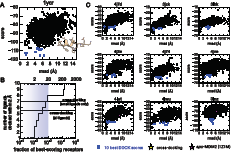
Scatter plots of DOCK scores versus the rmsd of the docked pose for all 2000 MSM receptor microstates show correlated funnel-like landscapes. (A) For the p53 ligand, the MSM receptor ensemble is more suitable for docking than any of the co-crystal receptor structures with other ligands. (B) The number of correct docking predictions found in some number of best-scoring poses (the true positive rate) for our test set is comparable to the cross-docking results. (C) Scatter plots for all ligands in our test set, shown with the 10 best-scoring poses docked to the MSM microstates (blue dots), cross-docking results (yellow stars), and docking results to the *apo*-MDM2 NMR structure (purple star, absences denote DOCK failures).

A potential caveat of these results is that the DOCK energy function is designed for the inexpensive evaluation of very large screening sets, at the potential cost of accuracy. For the PMI N8A peptide ligand, the lowest-energy DOCK score consistently predicts a non-native pose in which the key tryptophan and phenylalanine are placed correctly in the binding site, but with non-native sidechain rotamers, tilting the PMI helix ~30° in the binding cleft. We explored several alternative protocols designed to test whether this was due to our artificial cyclization scheme used to fix the backbone, or other search parameters; based on similar results in all cases, we conclude that scoring function accuracy is responsible.

### Simulation of functional lid motions is key to successful computational docking

Since our simulations started from an *apo*-MDM2 structure with a closed binding cleft not amenable to computational docking, we were curious to see how the functional lid motions identified in the MSM might be related to the generation of docking-competent receptor structures. A projection of the DOCK scores to the tICA landscape reveals that a significant clustering of low-scoring poses are found on the far right edge of the landscape, corresponding to states where the lid region is associated with the binding cleft (Figure 5a). This feature is more pronounced for the peptide docking results, but can also be seen clearly for the small-docking results (Figure S6). In previous work, we performed a number of *apo*-MDM2 simulations in various force fields, with trajectory lengths up to 1 *μ*s. The projection of these data onto the tICA landscapes shows that, regardless of the force field chosen, these simulations do not sample the full extent of lid motion seen in the MSM (Figure 5b).

**Figure 5.**
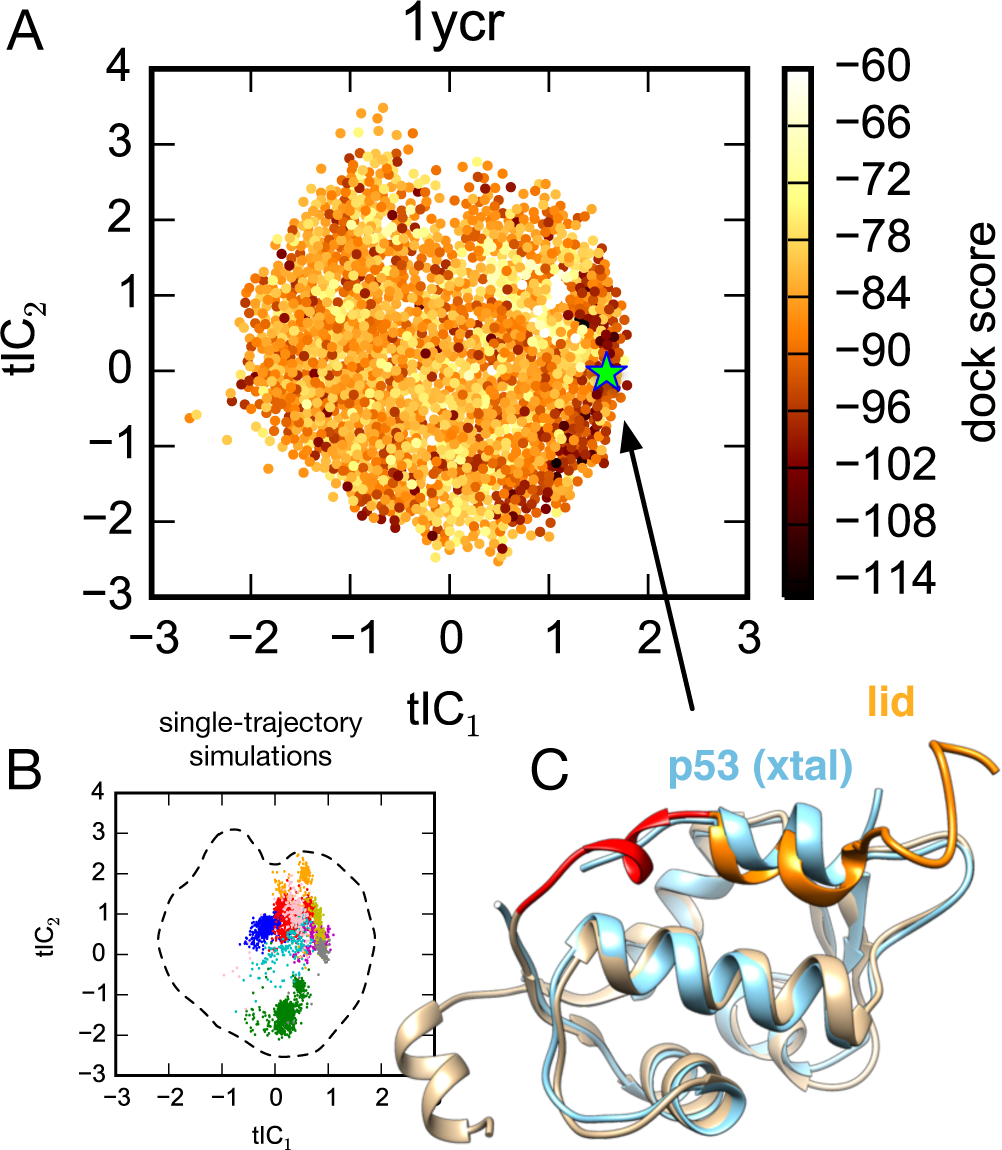
(A) Projection of the DOCK scores for p53+MDM2 to the tICA landscape reveals a significant clustering of low-scoring poses corresponding to lid-associated structures. (B) Previous 200-ns and 1-*μ*s sing<u>l</u>e-trajectory simulations of *apo*-MDM2 by Pantelopulos et al.^32^ projected to the tICA landscape show that simulations do not sample the full extent of lid motion seen in the MSM. Simulations were performed in force fields AMBER ff14sb (1 *μ*, blue), ff99sb-ildn-nmr (1 *μ*s, red), ff99sb-ildn (200 ns, cyan), ff99sb (200 ns, yellow), ff99sb-ildn-phi (200 ns, orange), ff14sb (200 ns, magenta), CHARMM22* (1 *μ*s, green), CHARMM36 (200 ns, pink). (C) The receptor structure with the lowest DOCK score (green star, panel A) exhibits a lid conformation closely mimicking the structure of p53 TAD bound to MDM2.

An inspection of the MDM2 receptor structures found on the far right of the tICA landscape, in the region of lowest DOCK scores, reveals many receptor conformations with their lid region associating with the MDM2 binding cleft. Indeed, the lowest-scoring receptor structure in this region for the p53 ligand (Figure 5a, green star) is revealed to have a helical conformation, closely mimicking the bound pose of the p53 transactivation domain (Figure 5c). In the unbound state, residues 11-17 (DGAVTTS) of the lid region have a low propensity to form a p53-like helix, forming helical structure when bound in the cleft (Figure S7).

## Discussion

Recent computational studies have examined how bound ligands and/or post-translational modifications modulate the conformational dynamics of the lid region.^33,34^ The results presented here complement this work, and strongly suggest that lid region association helps to induce binding-competent, open-cleft receptor structures amenable to computational docking. This idea is consistent with the induced-fit “fly-casting” mechanism that has been proposed as the dominant mechanism of many intrinsically disordered peptides that fold upon binding,^35^ including the p53 TAD of MDM2.^17^ Previous 200-ns and 1-*μ*s simulations of *apo*-MDM2 starting from an initial closed-cleft NMR structure sample a range of open-and closed-cleft structures, but do not visit receptor structures highly competent for p53 binding, presumably because in these trajectories the lid region doesn’t sufficiently associate with the cleft to induce such structures. These findings underscore the utility of large-scale conformational sampling and analysis made possible by Markov State Model approaches. Indeed, the large-scale MD+MSM approach we use here is able to identify MDM2 receptor structures as good or better than available crystal structures for computational docking. In the future, MSMs are likely to be a valuable component of emerging molecular simulation-based methods for ensemble-based virtual screening,^36^^−^^38^ especially for homology models.^39^

Given the known limitations in the accuracy of scoring functions for computational docking, we expect that the use of MD+MSM simulated receptor ensembles will perform even better in conjunction with more accurate energy functions, especially as a starting point for more sophisticated methods such as free energy perturbation,^32^ for which elucidation of relative binding modes is especially important.^40^

Finally, we note that many drug targets are cell signaling proteins regulated in some way by intrinsically disordered binding partners. Many of these also have intrinsically disordered auto-inhibitory sequences than can mimic these natural substrates. For example, p53 binding partner MDMX was recently found to have an auto-inhibitory domain that inhibits binding through structural mimicry of the p53-MDMX interaction,^41^ a discovery which helps explain the failure of prior small-molecule drug screening efforts that did not utilize the full-length target. Similarly, our results suggest that explicit consideration of such disordered regions in simulation models may be much more important than currently appreciated, and could lead to greater functional insights and more successful computational drug discovery efforts.

## Conclusion

Large-scale molecular simulation combined with Markov State Model analysis of simulated *apo*-MDM2 dynamics predicts diffuse, yet two-state binding of its disordered lid region to the p53 cleft, consistent with experiment. Computational docking of known MDM2 ligands to this simulated receptor ensemble achieves success comparable to crystal structure cross-docking, suggesting that virtual screening studies can benefit from Markov State Model approaches. These results underscore the importance of the disordered lid region in both understanding MDM2 functional motions and in computational drug discovery.

## Methods

### Molecular Simulation

GROMACS 4.5 was used for all simulation preparation and production.^42^ Twenty-four initial conformations of the p53-binding region of *apo*-MDM2 (residues 1-119) were taken from the NMR-derived structure (PDB: 1Z1M).^43^ The AMBER ff99sb-ildn-nmr force field^44^ was chosen based on previous work demonstrating its accuracy and ability to predict initial structuring of the lid region in 1 *μ*s simulations.^32^ All systems were constructed as periodic cubic boxes solvated with 17268 explicit TIP3P waters and 0.1 M NaCl. Stochastic (Langevin) dynamics was simulated using a leap-frog integrator with a time step of 2 fs and an inverse friction constant of 1 ps. Non-bonded cutoffs of 0.9 nm were used for both real-space Particle-Mesh Ewald (PME) electrostatic and vdW interactions. Protein and non-protein atoms were temperature-and pressure-coupled as separate groups in the Berendsen thermostat, at 300K and 1 atm, using a 1 ps time constant, compressibility of 4.5×10^−5^ bar^−1^. Prior to production runs, all systems were equilibrated in the isothermal-isobaric (NPT) ensemble until the system volume converged to 538.71 nm^3^. Pro-duction runs in the canonical (NVT) ensemble were performed on the Foldinghome distributed computing network,^45^ obtaining 175.7 *μ*s of aggregate trajectory data. The distribution of trajectory lengths is roughly exponential, with a maximum trajectory length of 945 ns, and average trajectory length of 67.0 ns (Figure S1).

### Markov State Model (MSM) construction

MSMBuilder^46^ was aused to construct MSMs from the trajectory data. Time-lagged independent component analysis (tICA) was performed using a tICA lag time of one snapshot (100 ps), to find a low-dimensional subspace best capturing the slowest motions of the N-terminus and its binding cleft. The subspace consists of linear combinations of the set of 2304 pairwise distances be-tween all C_α_ atoms either in residues 1-24 of MDM2, or within 5 Åof any atom of the p53 helix in the crystal structure of *holo*-MDM2 (PDB: 1YCR). Conformational clustering in this low-dimensional subspace was used to define a set of 2000 metastable microstates. A generalized matrix Rayleigh quotient (GMRQ) method^47^ was used to find optimal MSM model parameters. This analysis, which involves a cross-validation procedure wherein the trajectory data is partitioned in testing and training sets, determined that (1) *k*-centers clustering produced marginally better models than k-means, (2) only two tICA components were needed to accurately capture the slowest conformational motions, (3) an MSM lag time of 100 ps produced the most accurate MSMs, and (4) the GMRQ score (reflecting model quality) plateaus around 2000 microstates (Figure S2). With metastable microstates suitably defined, the matrix of transition prob-abilities 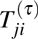 of transitioning from state *i* to state *j* within lag time τ was computed using a maximum-likelihood estimator from the observed transition counts.^46^ Coarse-graining of MSM microstates into a 150-macrostate model was performed using the BACE algorithm.^48^

### Structural analysis

Analysis of trajectory data was performed using the MDTraj python library. Secondary structure populations were com-puted using the DSSP algorithm, with helical states corresponding to DSSP assignments G, H, I; sheet states corresponded to DSSP assignments B and E. The SHIFTX2 algorithm^29^ was used to predict chemical shift values, using 10x subsampling of trajectory snapshots, for each MSM macrostate. To quantify the significance of interresidue contacts formed in specific conformational states, we compute a Bayes Factor (BF) contact metric for each residue pair in MDM2^25^. More details about this are given in the Supporting Information.

### Computational docking with DOCK6

Computational docking was performed using UCSF DOCK version 6.7.^41^^−^^51^ The crystal structure coordinates were downloaded from the PDB and processed using the UCSF Chimera dockprep tool.^51,52^ Small molecules were assigned AM1-BCC ligand partial charges with AmberTools antechamber,^53^ while peptide ligands were assigned ff14SB charges. Frames taken from each of the 2000 microstate clusters were converted into DOCK-compatible MOL2 files. Owing to inconsistencies in hydrogen atom naming schemes, each such frame was reassigned optimized instantaneous protonation states using the REDUCE tool.^54^ Grids at 0.3 Å-resolution were computed for each of the 2000 MD-derived frames. In order to improve sampling, each rigid segment with five or more atoms (e.g. pyrroles or larger) was used as an anchor during small molecule docking. A unique feature of the DOCK program is the anchor and grow algorithm.^49^ A rigid section of the molecule, often a large aromatic scaffold (anchor) is first oriented in the binding site. The remaining torsions are then grown one-by-one, clustering and pruning unfavorable conformations at every step until a final set of viable fully grown conformers remain. This breadth-first search approach takes exponential computational time, which severely limits docking of larger molecules. DOCK 5 was only tested on a set of molecules with seven or fewer rotatable bonds.^55^ For DOCK 6.2 onwards, the addition of a fast internal energy score, coupled with aggressive pruning and rmsd symmetry, allowed reasonable performance with larger molecules (65.5% success on 8-15 torsions and 48% with >15 torsions).^49^ Earlier work^49^ demonstrated that despite these gains, docking success drops linearly with the number of rotatable bonds, while runtime increases exponentially. DOCK considers closed cycles in molecules to be rigid when sampling torsions. However, the simplex minimizer still relaxes local backbone conformations within these cycles. Thus, for peptide ligands, we introduced an artificial bond between the N-and C-termini to ‘rigidify’ the backbone for the purposes of docking. This ameliorates the need to fold alpha helical ligands *ab initio* with the limited molecular mechanics scoring function van der Waals and electrostatics with a distance dependent dielectric) in DOCK. In the case of the p53 TAD fragment, this reduces 66 torsions to 29 torsions after rigidifying the backbone. DOCK thus considers the backbone to be an anchor, with each sidechain torsion grown *in situ* for each receptor microstate. Cases where the receptor conformation does not (1) accommodate the backbone, or (2) allow all the sidechains to complete growth, forces the docked ligand out of the binding site, resulting in a poor interaction score.

## Acknowledgements

This research was supported in part by the National Science Founda-tion through major research instrumentation grant number CNS-09-58854, and MCB-1412508. G.A.P. is supported by a National Science Foundation Graduate Research Fellowship under Grant DGE-1545957.

### Author contributions statement

V.V. conceived the experiments, G.P. and S.M. conducted the experiments. All authors analyzed the results and reviewed the manuscript.

### Additional information

Competing financial interests. No competing financial interests exist for any of the authors.

## Supplementary Information

Markov models of the *apo*-MDM2 lid region reveal diffuse yet two-state binding dynamics and receptor poses for computational docking Sudipto Mukherjee, George A. Pantelopulos and Vincent A. Voelz Department of Chemistry, Temple University, Philadelphia, PA

**Supporting Text**

**Generalized Matrix Raleigh Quotient (GMRQ) Analysis**

To choose optimal parameters for MSM construction, we use the GMRQ method recently developed by McGibbon et al.^1^ This method exploits the variational principle of conformational dynamics,^56^^−^^58^, that any approximation to the true eigenvectors of a dynamical operator will necessarily underestimate its eigenvalues, which correspond to the relaxation timescales. For an MSM of *n*discrete conformational states indexed by *k* = 1…*n*, approximations to the eigenfunctions are linear combinations of indicator basis set functions *ϕ*_*k*_(*x*) (equal to 1 if *x* is in state *k* and 0 otherwise.) The generalized matrix Raleigh quotient R quantifies how well a given linear combination of basis functions captures the *m* slowest eigenvectors/timescales. It is computed as

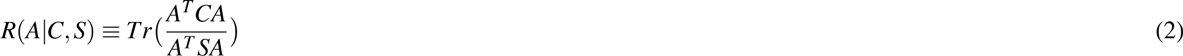
where *A* is an *n* × *m* matrix whose *j*^*th*^ column contains the linear coefficients to approximate the *j*^*th*^ c eigenvector as ∑_*k*_*A*_*kj*_*ϕ*_*k*_(*x*), *C* is an *n* × *n*time-lagged correlation matrix of the indicator basis functions calculated from the trajectory data, and *S* is an *n* x *n* covariance matrix of the indicator basis functions. For an MSM, both *C* and *S* can be estimated from the number of transitions between states observed in the simulation trajectories. Finding the coefficient matrix *A* that maximizes *R* is the same eigenproblem solved in the tICA method.^2,3^ Once maximized, *R* can be used to evaluate the quality of the MSM state decomposition by how well it captures the true conformational dynamics.

To avoid overfitting to the data, a cross-validation approach is used in which the simulation data is partitioned into 5-fold leave-one-out training/testing sets. In five separate trials, *R* is maximized using 4/5 of the trajectory data, which we report as the GMRQ training score. Using these optimal values of the coefficients in *A*, the remaining 1/5 of the data is then used to compute *R*, which we report as the GMRQ testing score.

To select the MSM that most accurately captures the conformational dynamics of the MDM2 lid region, we explored various model construction parameters and chose the model with the largest GMRQ testing score (Figure S2). The time step between collected trajectory snapshots is *τ*_0_ = 100 ps. We explored MSMs built using different tICA lag times (i.e. the lag time to calculate the distance correlation matrix needed to find the tICA components) and different MSM lag times. For MSMs constructed using a range of 200 to 2000 microstates, a lag time of 1 (in units of *τ*_0_) is optimal (Figure S2a,b). We also explored MSMs constructed using different numbers of tICA components (2 to 15 tICs) as the subspace to project trajectory data and perform conformational clustering. We found that, for MSMs constructed using a range of 200 to 2000 microstates, projections utilizing 2 tICs give the best GMRQ testing scores (Figure S2c). In all of our tests, GMRQ testing scores versus the number of MSM microstates plateau near 2000 microstates, so we chose this number of microstates for MSM construction.

Given that typical MSM lag times for protein folding and binding studies range from 1-200 ns, it is somewhat surprising that such a short lagtime (100 ps) is preferred for MSM construction. To validate this result, we additionally built MSMs using lag times of 2, 5, 10, 100 and 1000 (in units of τ_0_). We find that the equilibrium populations and slowest relaxation mode eigenvector *ϕ*_1_ are remarkably robust at all lag times, with the exception of 1000, for which finite sampling artifacts become pronounced (Figure S3a). A likely explanation for the short lag time being optimal is the highly diffusive nature of the lid region dynamics, coupled with a trajectory data set enriched in many short simulations (see Figure S1). To test this idea, we built MSMs using lag times of 1, 10, 100 and 1000 using the same construction parameters, but without using a standard ergodic trimming step, which is usually employed to avoid statistical bias from non-equilibrium trajectories.^4^ This bias is particularly pronounced for MSMs built from distributed computing simulations, due to the use of many short trajectories that make forward transitions to new states, but no backward transitions. Enforcing detailed balance on a MSM built from this data can therefore introduce “trap” artifacts in which states can have incorrectly high population estimates. Indeed, without ergodic trimming, the tICA model strongly exhibits the presence of traps, indicating the non-ergodicity of the underlying trajectory data arising from diffusivity and trajectory length (Figure S3b).

**Bayes Factor analysis of inter-residue contacts**

To quantify the significance of inter-residue contacts formed in specific conformational states, we compute a Bayes Factor (*BF*) contact metric for each residue pair in MDM2.^5^ The *BF*_*k*_(*i*, *j*) for contacts between residues *i* and *j*, given the protein is in some conformational state *k*,s is computed as

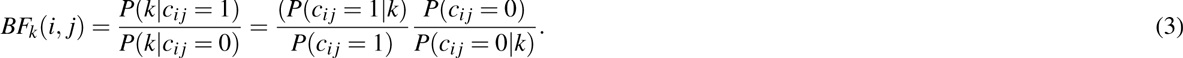

Here, *c*_*ij*_ is an indicator variable that takes the value of 1 if a contact between residues *i* and *j* is present, and 0 otherwise. The *BF*_*k*_(*i,j*) value can be thought as the statistical over-representation of contact *c*_*ij*_ in conformational state *k*, and hence a measure of its importance in uniquely defining the structural features of that state. For example, if *BF*_*k*_ = 2, that means that the equilibrium constant for contact formation between residues *i* and *j* is twice as large for state *k* as it is for the whole ensemble. We compute Bayes factors for contacts separated by three or more residues in sequence, and define a contact formed between two residues if any two non-hydrogen atoms are closer than 4 Å.

**Supporting Figures**

Figure S1. Trajectory length distributions Figure S2. GMRQ Analysis Figure S3. Effects of lag time and/or ergodic trimming on constructed MSMs Figure S4. Changes in inter-residue contacts and secondary structure along eigenmode relaxations *ϕ*1 and *ϕ*2. Figure S5. Bayes Factors of inter-residue contacts for tICA landscape quadrants. Figure S6. DOCK scores projected onto the 2D tICA landscape for all ligands Figure S7. Backbone RMSDs of MDM2 lid residues 11-17 to bound-state p53 helix. Supporting References

**Figure S1.**
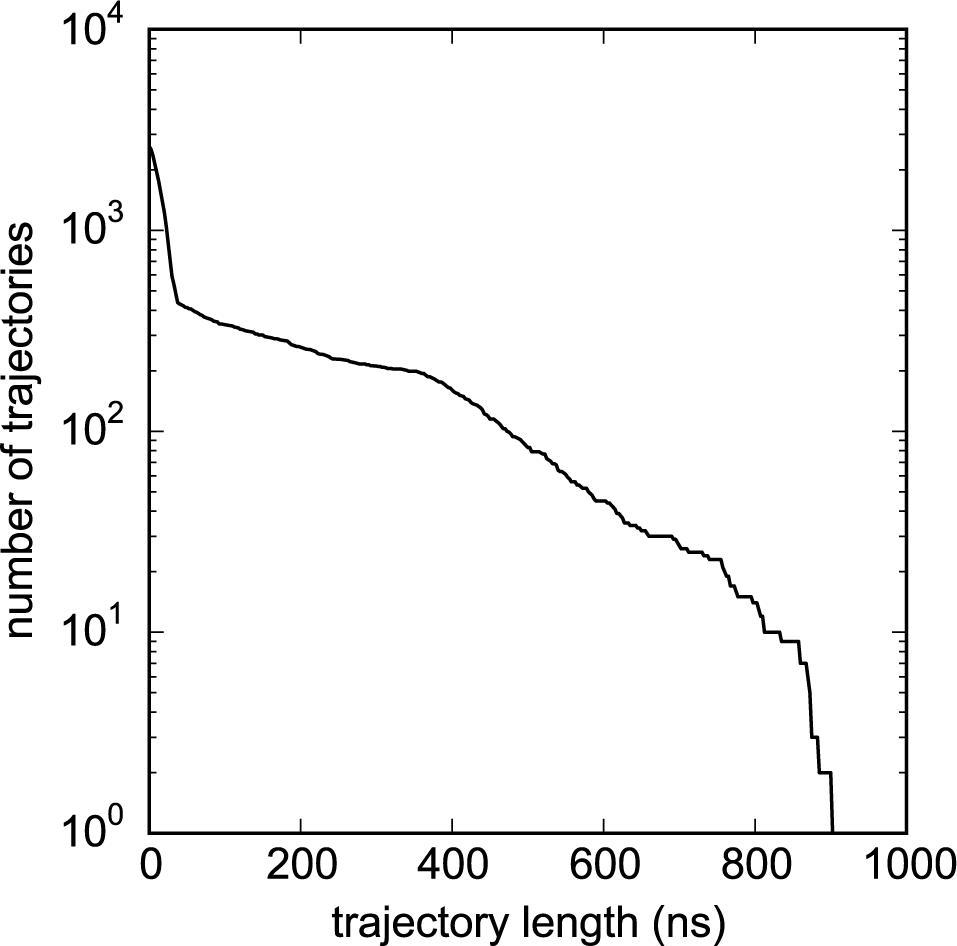
Distribution of trajectory lengths for the 175.7 *μ*s of aggregate trajectory data simulated on the Foldinghome distributed computing network. The maximum trajectory length is 945 ns, and average trajectory length is 67.0 ns.

**Figure S2.**
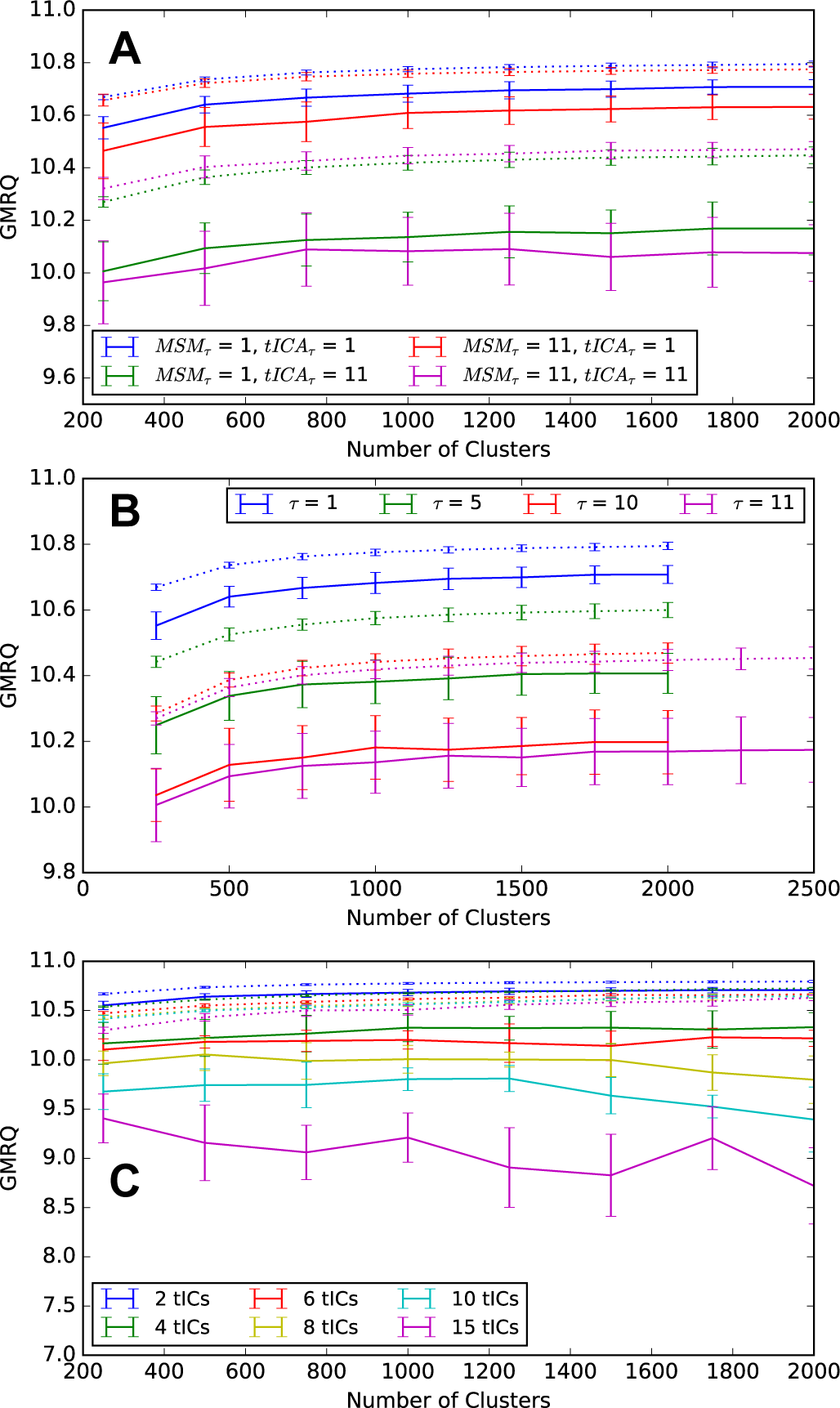
Mean GMRQ training scores (dotted line) and testing scores (solid line) computed using 5-fold cross validation, shown as a function of the number of *k*-centers clusters used to build the MSM. (A) GMRQ scores calculated using slow (11τ_0_) and fast (1τ_0_) MSM and tICA lag times. (B) GMRQ scores calculated using a range (2τ_0_ to 11τ_0_) of MSM lag times; each MSM was constructed using clustering in the subspace defined by the 2 largest tICA components. (C) GMRQ scores calculated for MSMs built from various numbers of tICA components (2 to 15). Error bars in all figures show standard deviations of the 5-fold cross-validation trials.

**Figure S3.**
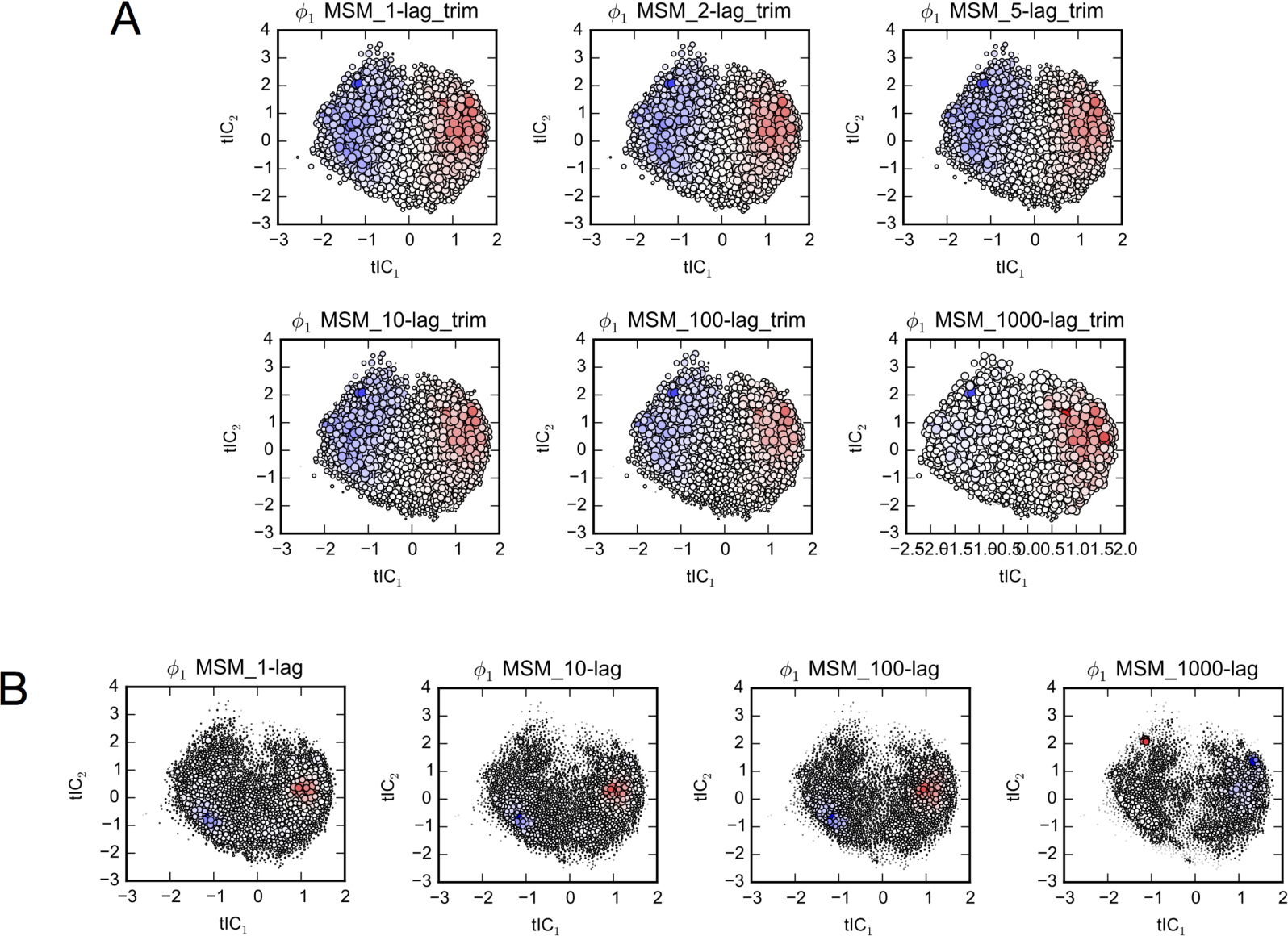
(A) Projections of the 2000 MSM microstates (filled circles) to tIC_1_ and tIC_2_ coordinates from MSMs built using lag times 1, 2, 5, 10, 100 and 1000 (in units *τ*_0_). The size of each circle is proportional to the logarithm of its equilibrium population, and is colored according to the slowest relaxation eigenmode, *ϕ*_1_. (B) MSMs built using lag times 1, 10, 100, 1000, without the standard ergodic trimming step.

**Figure S4.**
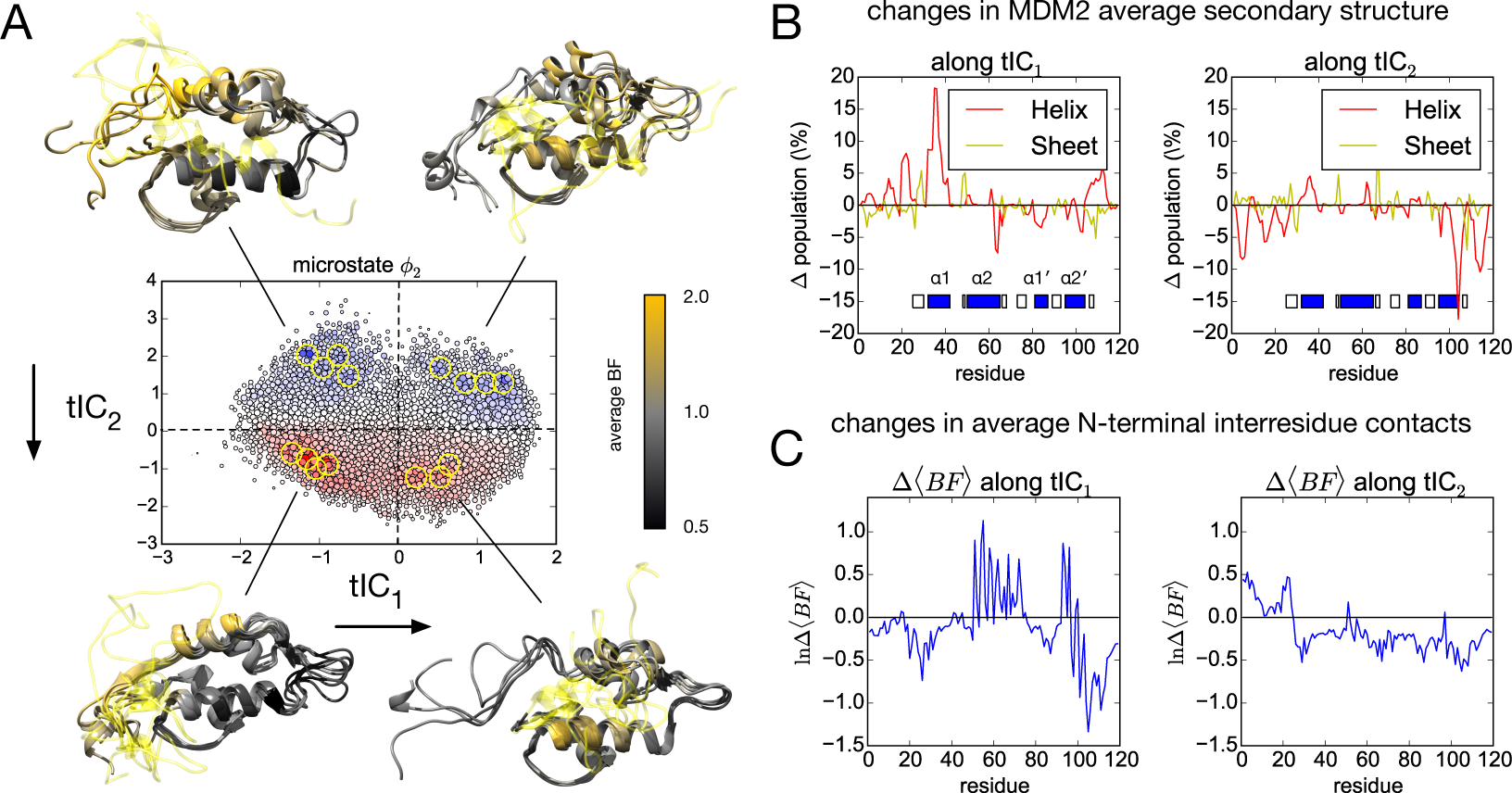
(A) Projection of the 2000 MSM microstates (filled circles) to tIC_1_ and tIC_2_ coordinates. The size of each circle is proportional to the logarithm of the equilibrium population, and is colored according to the second slowest relaxation eigenmode, *ϕ*_2_. Population flux along this mode is from blue to red, predominantly representing the structuring of the MDM2 termini, which we visualize using five representative structures from each basin (circled in yellow on the tICA landscape). Average Bayes Factors for contacts between lid region (yellow ribbon) and non-lid residues are shown as a color gradient (black to orange) on the ribbon structure of MDM2. (B) Differences in per-residue helix and sheet content of MDM2 are shown along tIC_1_ (i.e. differences between conformations with positive vs. negative values of tIC_1_) and along tIC_2_. (C) Changes in the ratio of mean Bayes Factors for contacts between the lid region (residues 1-25 of MDM2) and all other residues, along tIC_1_ and tIC_2_.

**Figure S5.**
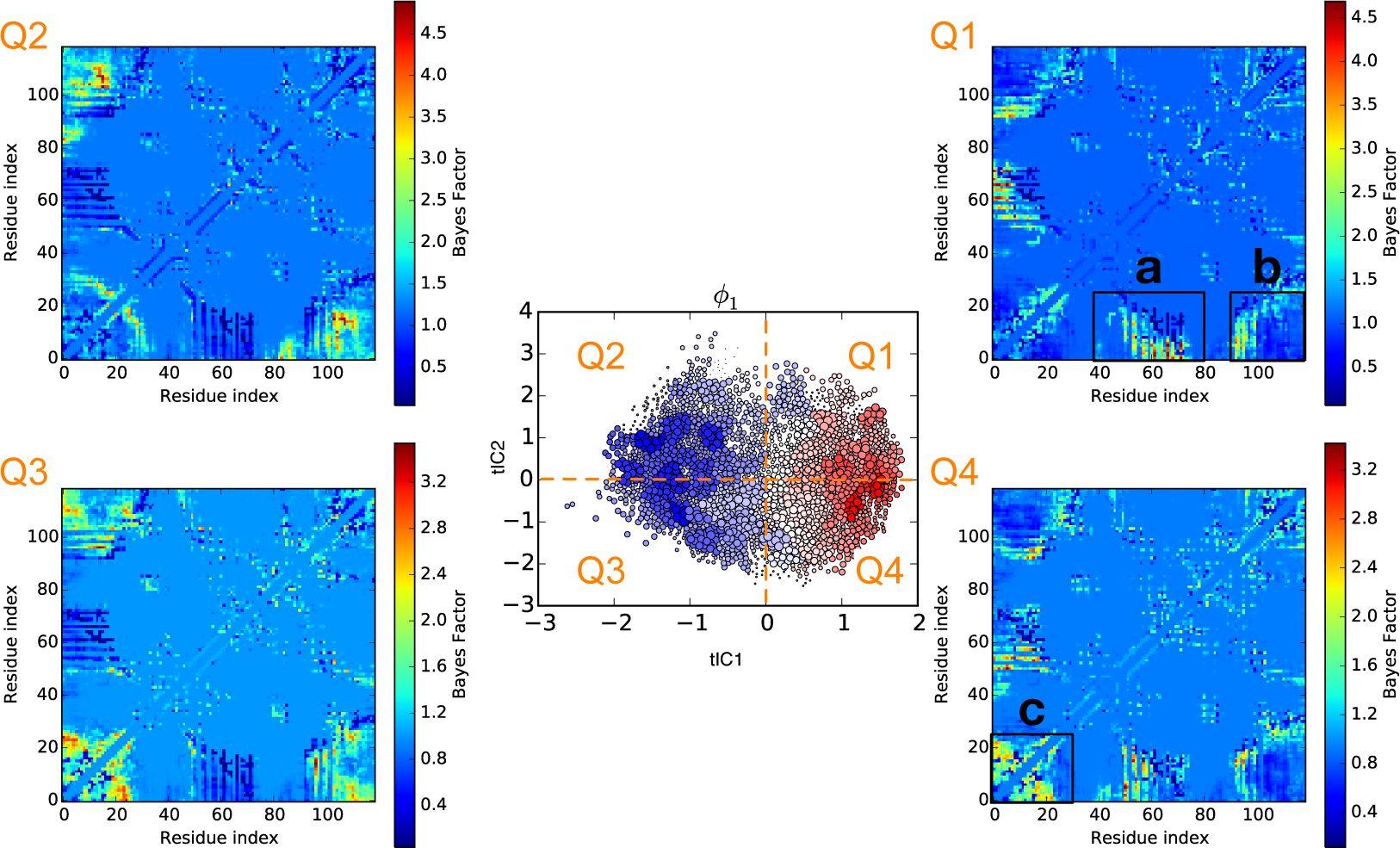
Bayes Factors of inter-residue contacts calculated for each quadrant in (tIC_1_, tIC_2_)-space. At center is shown the 2000 MSM microstates projected to the tICA landscape, divided into quadrants Q1, Q2, Q3 and Q4. (a) Differences in the Bayes factors for Q2+Q3 versus Q1+Q4 reveal that contacts are formed between the N-terminus and the active site helix α_2_ along tIC_1_, the slowest relaxation mode, and (b) contacts are lost between the N-and C-terminus along tIC_1_. Differences in the Bayes factors for Q1+Q2 versus Q3+Q4 reveal that (c) conformational changes along tIC2 correspond to structuring in the N-terminus.

**Figure S6.**
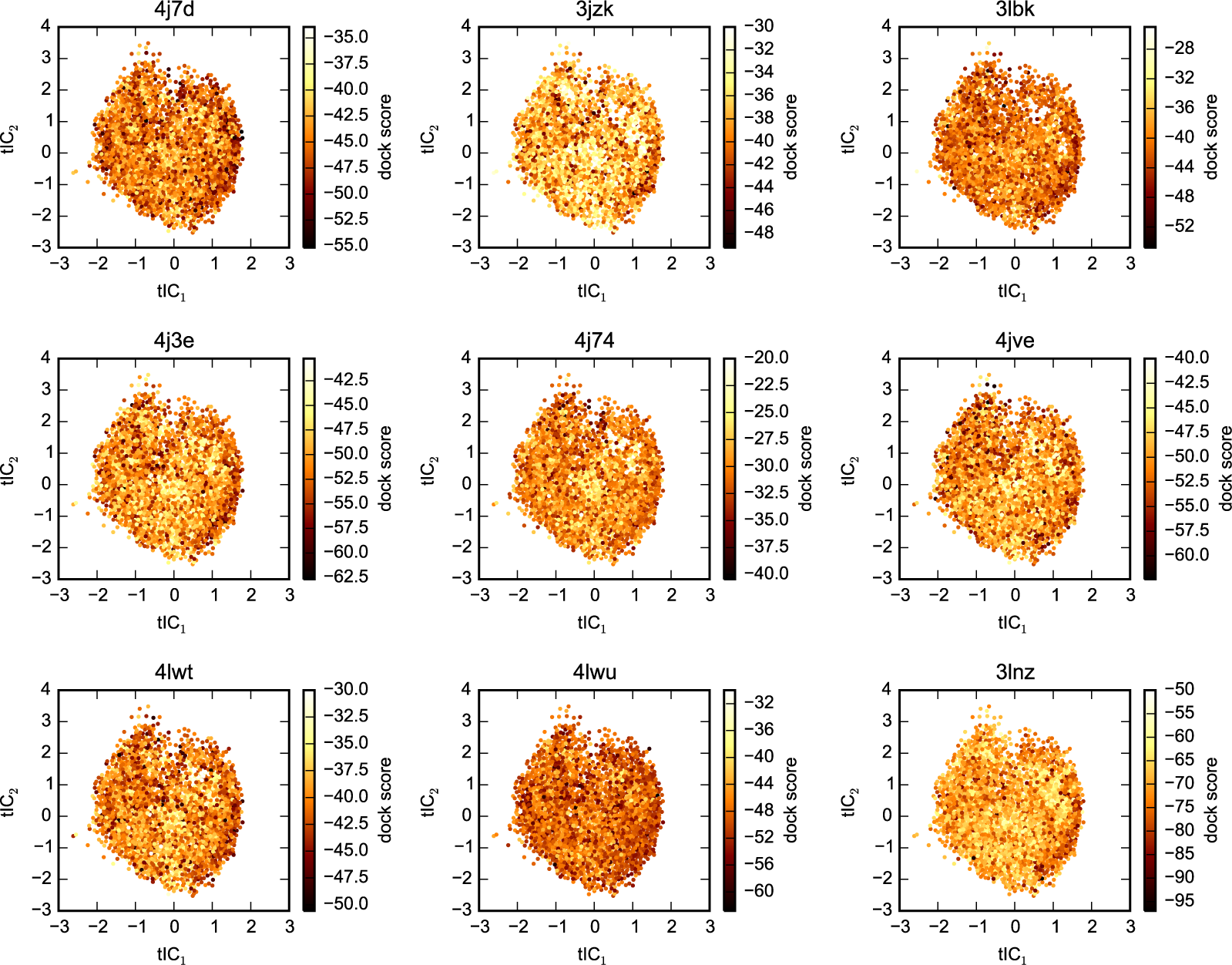
DOCK scores projected onto the 2D tICA landscape for all ligands. Results for p53 (1ycr) are shown in the main text.

**Figure S7.**
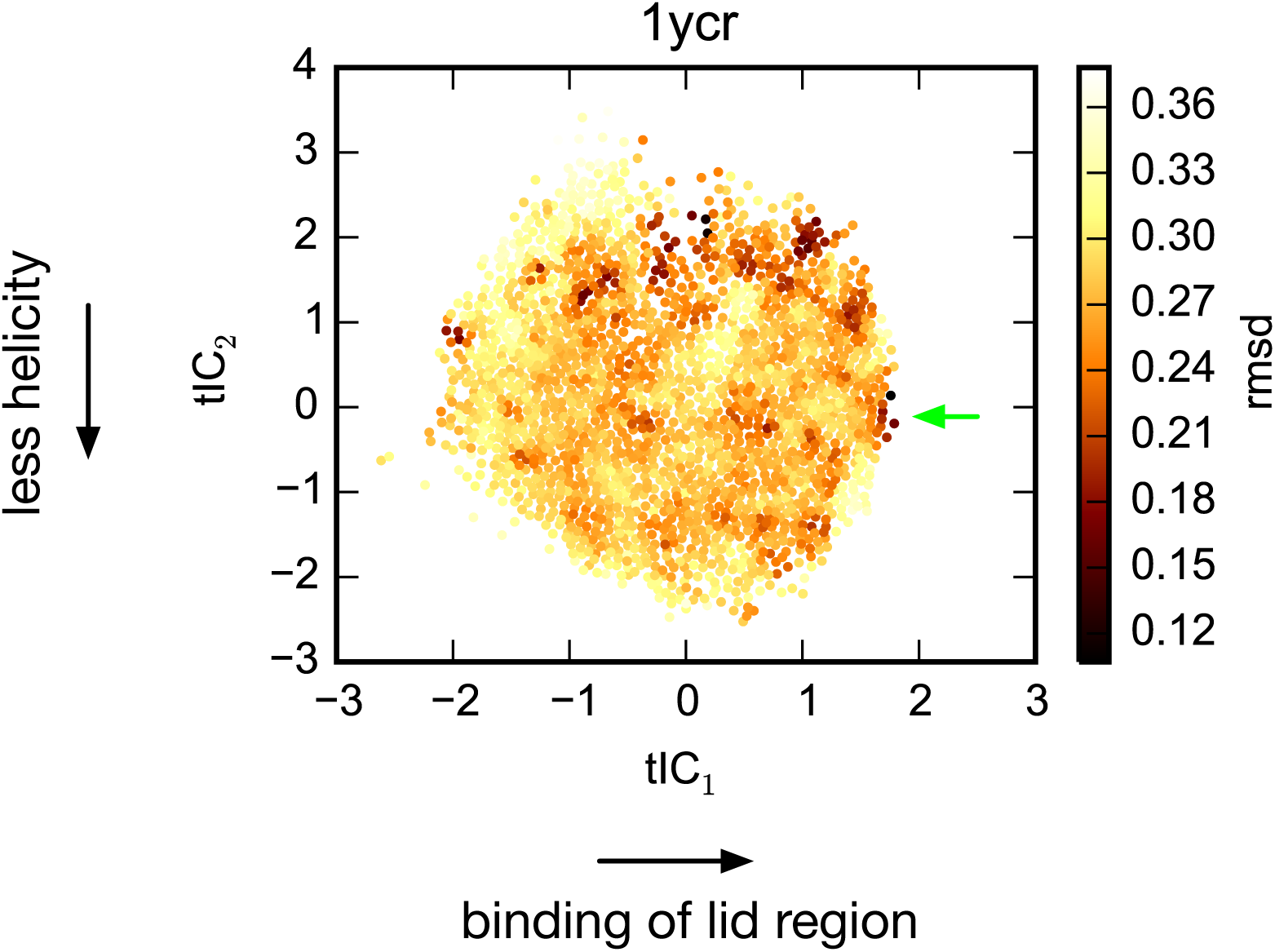
Backbone RMSDs of MDM2 lid residues 11-17 (DGAVTTS) to the bound-state p53 helix, shown for all 2000 MSM microstates on the tICA landscape. In the unbound state, this sequence has low propensity to form a p53-like helix, with two exceptions: (1) There is some residual helicity in the unbound state, which is lost along the *ϕ*_2_ eigenmode relaxation (positive to negative tIC_2_ values, see Figure S4) as the N-terminus self-associates, and (2) association of the lid region with the p53 binding cleft induces structuring of residues 11-17 to a conformation very similar to bound p53 (green arrow).

